# Monitoring of vaccine targets and interventions using global genome data: vaccines.watch

**DOI:** 10.1101/2025.06.13.659488

**Authors:** Sophia David, Khalil Abudahab, Natacha Couto, Wa Ode Dwi Daningrat, Corin Yeats, Alison Molloy, Philip M Ashton, Nabil-Fareed Alikhan, David M Aanensen, NIHR Global Health Research Unit on Genomics and enabling data for Surveillance of AMR

## Abstract

The expansion of pathogen genome sequencing into routine disease surveillance programmes is set to bring rapidly-growing volumes of increasingly structured data on a global scale. This has the potential to deliver exciting opportunities for accelerating vaccine development and monitoring. Here we present an interactive platform, vaccines.watch (https://vaccines.watch), which aims to support decision-making around vaccine formulations and roll-out by enabling interrogation of vaccine target diversity from global genome data. We have initially focused on targets included in existing or prospective multivalent polysaccharide-based vaccines for *Streptococcus pneumoniae*, *Klebsiella pneumoniae* (and related species) and *Acinetobacter baumannii*. The platform currently displays data for >100k high-quality genomes with geotemporal sampling information (post-2010), with new genomes assembled, analysed and incorporated on an ongoing basis (every 4 hours) as public data are newly deposited. Crucially, users can view vaccine target information in the broader context of genotypic variants (e.g. sequence types) and antimicrobial resistance markers. The platform also enables users to review the composite serotypes of pneumococcal vaccine formulations among the available genomes. For example, using data in vaccines.watch from 3 June 2025, we observed that serotypes included in the PCV13 and PCV21 formulations accounted for 36.2% (11,907/32,918) and 87.4% (28,764/32,918) of global public genomes, respectively. The platform also enables continuous review of the global genomic landscape of the included pathogens, enabling identification of gaps (e.g. in geographic coverage) that should be targeted with increased genomic surveillance. Indeed we demonstrate that substantial geographic gaps remain in the coverage of available genomes, with over half of countries contributing no genomes for each of the three pathogens. However, while caution in interpretation is important, as global representativeness of genome data grows, vaccines.watch is positioned to support different stages of the vaccine pipeline, from selection of target antigens to post-rollout monitoring of population changes.

## Introduction

Vaccines have been a highly effective intervention for decreasing the burden of infectious diseases. They also have the potential to play an important role in reducing antimicrobial resistance (AMR), both by preventing infections caused by antimicrobial-resistant (and - susceptible) strains and by decreasing antimicrobial usage [1]. This has been exemplified by the recently-introduced typhoid conjugate vaccine (TCV) in Pakistan, which has been shown to be over 90% effective in protecting young children against extensively drug-resistant (XDR) typhoid infections [2]. A major advantage of vaccines is that they typically have a sustainable impact in reducing disease burden, with resistance to vaccines evolving far less readily than antimicrobial resistance [3]. Yet despite the success of vaccines, we still do not have any available vaccines for many major pathogens, while others necessitate improved or updated formulations [4]. Recently, the World Health Organisation (WHO) identified 17 endemic pathogens, including several bacterial species with a high AMR burden, for which global efforts in vaccine research and development should be prioritised [5].

Genomic information has been used in vaccine development since the availability of the first bacterial genomes, in particular by enabling high-throughput *in silico* screening of all protein-encoding genes to identify potential target antigens. This approach, known as reverse vaccinology [6], has been used in the development of highly effective vaccines against *Neisseria meningitidis* serogroup B (MenB) [7, 8] as well as in antigen selection for multiple other vaccines since [9, 10]. An increasing array of bioinformatic tools have also been developed to further aid antigen selection, which make use of the growing biological databases (including genomic, transcriptomic and proteomic data) and increasingly sophisticated analytical and machine-learning methods [11]. In particular, pan-genome methods are typically used to identify genes that are common (core) to all strains of the target pathogen, as well as tools that predict protein characteristics (e.g. immunogenicity, subcellular localisation), epitopes, and the potential toxicity and allergenicity of candidate antigens.

Over the coming years, the expansion of pathogen genome sequencing into routine disease surveillance programmes is set to bring rapidly-growing volumes of increasingly structured data on a global scale. This ongoing shift towards widespread genomic capacity, supported by major international public health agencies including the WHO [12], will provide increasing opportunities to accelerate vaccine development using genomic data. In particular, detailed genomic insights into contemporary pathogen population dynamics gained from surveillance data will enable us to better optimise vaccine formulations (including those targeted at specific populations or regions), improve our understanding of the impact of vaccine rollout on pathogen diversity, and more precisely monitor and deduce mechanisms of vaccine escape.

However, technical barriers to fully capitalising on the available and upcoming opportunities remain, both around accessing the growing volumes of genomic data from the public sequence archives, and processing and translating the data into relevant insights across different pathogens for relevant stakeholders. We recently described our amr.watch platform (https://amr.watch) that analyses and visualises AMR trends from global public genome data [13]. For each of the priority bacterial pathogens defined by the WHO [14], amr.watch incorporates and displays data on an ongoing (“always-on”) basis using relevant analytics, with the aim of supporting both research and policy. The platform also enables continuous review of the global landscape of pathogen genome sequencing, revealing opportunities whereby genomic data may already be able to reliably inform public health interventions as well as identifying gaps (e.g. in geographic coverage) that can be targeted with increased surveillance efforts.

Using an extension of the “always-on” genome retrieval and analytical pipeline from amr.watch, here we present a sister platform, vaccines.watch (https://vaccines.watch), an interactive web application for assessing the prevalence and distribution of key vaccine targets from global genome data. We initially focus on providing support for the development and monitoring of multivalent vaccines with polysaccharide targets, the high diversity of which can necessitate choices over the particular polysaccharide types to include within a vaccine. Currently, vaccines.watch displays data on *Streptococcus pneumoniae* capsular-based serotypes, which form the targets of all licensed pneumococcal vaccines [15]. It also provides data on the capsule (K) and lipopolysaccharide (LPS) O-antigen (O) types from *Klebsiella pneumoniae* (and related species), as well as the capsule (K) and lipooligosaccharide (LOS) outer core (OC) antigen types from *Acinetobacter baumannii*. These are targets for novel vaccines, monoclonal antibody and phage therapies in both pathogens [16–19]. The vaccines.watch platform aims to provide scientists and vaccine developers with additional insights into the pathogen population dynamics around key vaccine targets, thereby supporting decision-making around vaccine formulations and roll-out.

## Methods

### Overview of the vaccines.watch platform

Vaccines.watch currently reports genome data from *S. pneumoniae*, *A. baumannii* and the *K. pneumoniae* species complex (SC). The latter includes five related species (see *Input data*). The platform uses data from genomes that are retrieved from the International Nucleotide Sequence Database Collaboration (INSDC) databases, curated and processed by Pathogenwatch (https://pathogen.watch) using a previously described workflow [13].

### Input data

The platform accepts data from paired-end Illumina genomes with a taxonomy ID recorded as 1313 (*S. pneumoniae*), 573 (*K. pneumoniae*), 1463165 (*K. quasipneumoniae*), 244366 (*K. variicola*), 2026240 (*K. quasivariicola*), 2489010 (*K. africana*) or 470 (*A. baumannii*) in the European Nucleotide Archive (ENA) metadata. We also require genomes to have an available sampling date from 2010 onwards, a sampling location that is decodeable to at least the country level, and a minimum of 20x coverage. We check for new genomes using the ENA Portal API every four hours and download genomes meeting the above criteria from the Sequence Read Archive (SRA) using the SRA-Toolkit fastq-dump v3.1.0 (https://hpc.nih.gov/apps/sratoolkit.html). The workflow has been described in detail previously [13], with specific filtering steps for the pathogens included here also provided at https://vaccines.watch/summary.

Sequence reads are assembled with a workflow (https://gitlab.com/cgps/ghru/pipelines/assembly) that uses the SPAdes assembler v3.15.3 [20]. We then assess the quality of the resulting assemblies with QUAST v5.0.2 [21] and verify the species using the Speciator tool (v4.0.0) (https://cgps.gitbook.io/pathogenwatch/technical-descriptions/species-assignment/speciator) within Pathogenwatch. For the *K. pneumoniae* SC, we accept genomes identified as any of the five species listed above, allowing for inconsistencies with the metadata due to known difficulties with phenotypic identification methods. Genomes that fail to meet quality criteria, as outlined for each pathogen in **Supplementary Table 1**, are excluded.

### Variant typing

The genotypic variants of the different pathogens are identified from the genome assemblies using community-based schemes implemented in Pathogenwatch. This variant typing comprises multi-locus sequence typing (MLST) for each of *S. pneumoniae* (PubMLST scheme), *K. pneumoniae* SC (BIGSdb-Pasteur scheme) and *A. baumannii* (Pasteur scheme) [22]. In addition to MLST, we also use Global Pneumococcal Sequencing Cluster (GPSC) assignments for *S. pneumoniae* [23] and “clonal group” assignments from the LIN code nomenclature for *K. pneumoniae* SC [24].

### Vaccine target typing

Kaptive v3.1.0 [25], implemented in Pathogenwatch, is used to identify the K and O loci (and predicted K and O types) from the *K. pneumoniae* SC genomes and the K and OC loci (and predicted K and OC types) from *A. baumannii* genomes (using the database described by [26] for the latter). For both pathogens, we only incorporate data into vaccines.watch from genomes where Kaptive has indicated that both the K and O/OC loci are “typeable”. SeroBA v2.0 [27], also implemented in Pathogenwatch, is used to identify the capsular polysaccharide (*cps*) loci and predict the resulting serotype from *S. pneumoniae* genomes. The Pathogenwatch implementation of SeroBA differs from the published method by using simulated reads generated from assemblies rather than the raw sequence reads, although inconsistencies between the methods occur rarely (0.14%) (see https://cgps.gitbook.io/pathogenwatch/technical-descriptions/typing-methods/seroba).

### Identification of mechanisms associated with AMR

For each pathogen, we identify genes and mutations associated with AMR for each of the antimicrobial classes reported in the WHO priority pathogens list [14]. These are identified using AMRFinderPlus v3.10.23, database version 2021-12-21.1 [28] with a curated database (**Supplementary Table 2**), as described previously [13].

### vaccines.watch web application

The development of vaccines.watch follows a similar framework to amr.watch, built using Next.js (https://nextjs.org/) and the React library (https://reactjs.org/). Geographic data are represented using Mapbox (https://www.mapbox.com) while the Apache ECharts library (https://echarts.apache.org/) is used for visualisation of data in charts (e.g. barplots). The vaccines.watch website incorporates the processed data from Pathogenwatch on an ongoing basis following successful implementation of the analytical steps described above. The *S. pneumoniae* vaccine formulations (up to 2025) available for selection within the vaccines.watch interface were obtained from a recent review [15]. All genomes and associated metadata visualised in vaccines.watch are also available within Pathogenwatch for further use by the community.

## Results

### Monitoring vaccine target diversity from global genomics data - the vaccines.watch platform

We have developed vaccines.watch (https://vaccines.watch), an interactive platform that enables monitoring of vaccine target antigens from pathogen genome data. We have initially focused on targets included in existing or prospective multivalent polysaccharide -based vaccines. Currently, these include capsular polysaccharide targets of *S. pneumoniae*, which form the basis of the licensed 23-valent pneumococcal polysaccharide vaccine (PPSV) and the different pneumococcal conjugate vaccine (PCV) formulations. We have also included the capsular and LPS O antigens of *K. pneumoniae* SC and the capsular and LOS OC antigens of *A. baumannii*, which are candidates for inclusion in novel vaccines and therapeutics in both pathogens.

As input, the vaccines.watch platform currently incorporates processed genome data from all high-quality short-read Illumina genomes in the INSDC databases from the relevant pathogens that have associated geotemporal sampling information and were sampled post-2010 (see *Methods*). Available public genomes are assembled and processed via Pathogenwatch on an ongoing basis (every 4h), with the resulting analytics displayed in vaccines.watch in real-time. This data includes phenotypic predictions of the target types generated from identification of the relevant genomic loci via the SeroBA [27] and Kaptive [25] tools. Additional information inferred from the genome assemblies, comprising the variant types (e.g. STs) and AMR mechanisms, are also incorporated. We provide a live overview of all genomes represented in vaccines.watch at https://vaccines.watch/all, while the filtering processes applied to the public data can be viewed at https://vaccines.watch/summary.

For each pathogen, vaccines.watch displays an interactive visualisation that enables rapid assessment and exploration of the diversity of vaccine target types and their geotemporal distribution (**Figure 1**). Vaccine target types are also placed into a broader population context by enabling assessment of their relationships with associated variant types (e.g. STs) and AMR markers. Users can survey the most frequent vaccine target types either globally, regionally or nationally, as well as assess specific target types of interest. In the case of *S. pneumoniae*, users can also select and explore specific sets of serotypes that are included in already-licensed pneumococcal vaccine formulations and vaccines under development. Visualisations with selected filters can be saved and/or shared onwards by users via the generation of URLs. All raw data shown in vaccines.watch can also be downloaded by users in CSV format.

**Figure 1.**
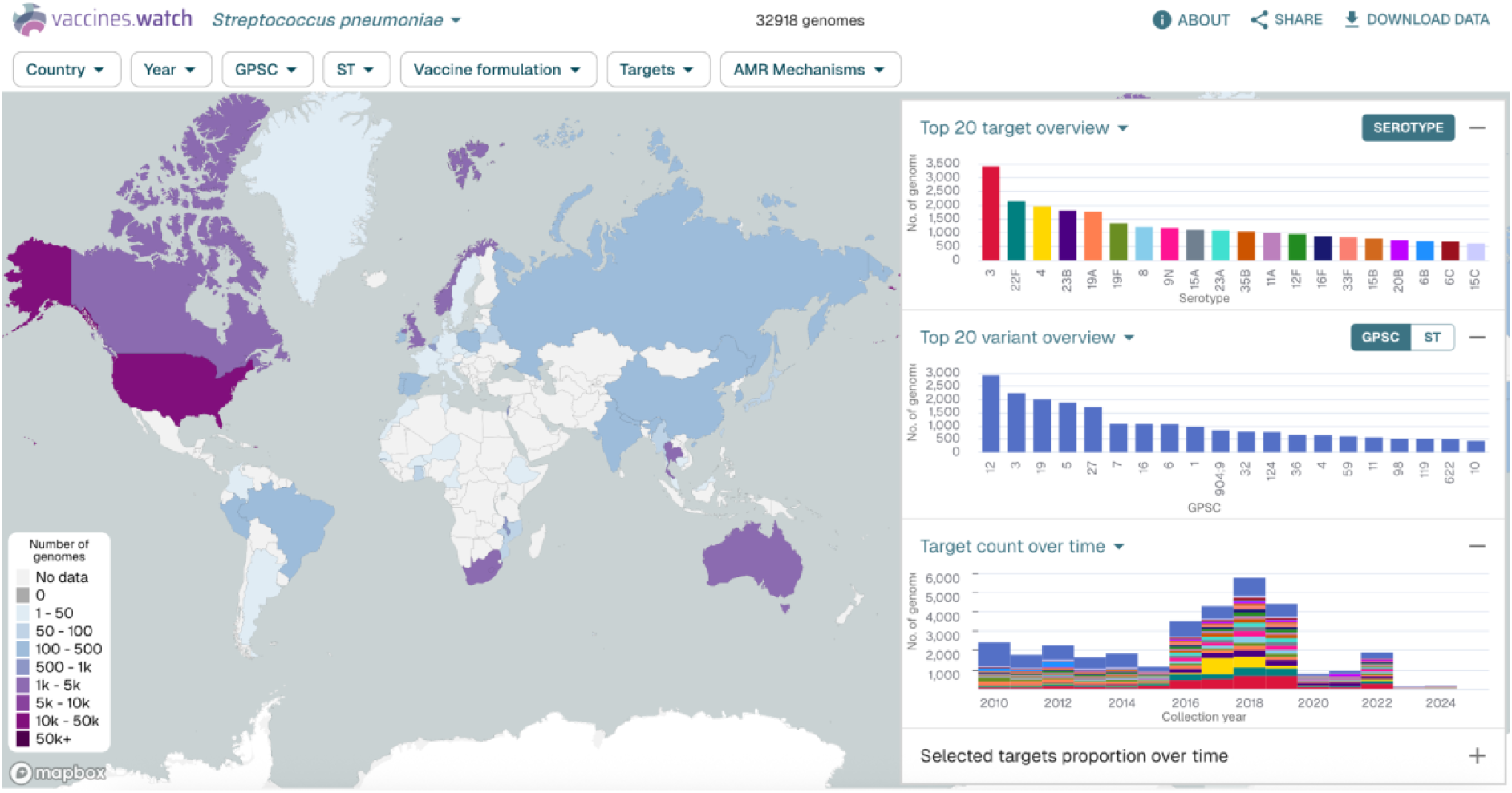
The interactive vaccines.watch platform here shows data for 32,918 *Streptococcus pneumoniae* genomes available as of 3 June 2025. The map shows the number of genomes sampled in each country. The right-hand figures show the twenty most frequent serotypes (top), the twenty most frequent variant types (global pneumococcal sequence clusters (GPSCs)) (middle) and the distribution of serotypes by sampling year (bottom). A similar visualisation, updated in real-time, can be found at: https://vaccines.watch/organism/1313

### Contemporary global landscape of *S. pneumoniae*, *K. pneumoniae* SC and *A. baumannii* genome sequencing

We recently reviewed the available public genome data displayed in our amr.watch platform for the WHO priority pathogens, which include the three pathogens included within vaccines.watch (except for the non-*K. pneumoniae* species in the *K. pneumoniae* SC) [13]. Here we provide an updated review for the pathogens included within vaccines.watch with additional detail. As of 3 June 2025, the vaccines.watch platform showed data for 32,918 *S. pneumoniae*, 49,035 *K. pneumoniae* SC, and 18,428 *A. baumannii* genomes from the public sequence archives (**Supplementary Table 3**). 94.1% (46,156/49,035) of the *K. pneumoniae* SC genomes belonged to the *K. pneumoniae* species, with a further 3.0% (1490/49,035) from *K. variicola*, 2.7% (1342/49,035) from *K. quasipneumoniae*, 0.07% (33/49,035) from *K. quasivariicola* and 0.03% (14/49,035) from *K. africana*. Notably, a large number of public genomes were excluded from the platform due to the absence of associated geographic and/or temporal metadata, including 110,361 *S. pneumoniae*, 39,031 *K. pneumoniae* SC and 12,154 *A. baumannii* (see https://vaccines.watch/summary for an updated summary). Among genomes that passed QC criteria and met metadata requirements, we also excluded a further 1576 belonging to the *K. pneumoniae* SC and 283 belonging to *A. baumannii* where the K and/or O/OC loci were identified as “untypeable” by Kaptive, ensuring the use of high-confident matches only.

For each of the three pathogens, we found that the number of genomes sampled from different geographic regions and individual countries was highly variable (**Figure 2**). Overall 79.2% (79,515/100,381) of all genomes were from high-income countries with 14.2% (14,217/100,381) from upper-middle, 5.1% (5154/100,381) from lower-middle and 1.5% (1470/100,381) from low-income countries. Almost half (47.1%; 15,519/32,918) of the *S. pneumoniae* genomes were from the USA, with 79.7% (26,250/32,918) of the total genomes from isolates sampled in seven countries (USA, South Africa, UK, Norway, Australia, Canada and Thailand). Approximately a third (32.0%; 15,681/49,035) of the *K. pneumoniae* SC genomes were from the USA with ten countries (USA, UK, China, Norway, Australia, Spain, Japan, Thailand, Italy and Germany) contributing 71.2% (34,899/49,035) of the total. *A. baumannii* genomes were the most unevenly distributed with approximately two-thirds from the USA (66.8%; 12,316/18,428) while China contributed an additional 10.3% (1902/18,428). Notably, we also identified substantial geographic gaps. In particular, over half of countries (55.8%; 139 of 249 with officially-assigned ISO-3166-1 codes) contributed no genome data for *K. pneumoniae* SC, which rose to 65.9% (164/249) for *A. baumannii* and 77.9% (194/249) for *S. pneumoniae*.

**Figure 2.**
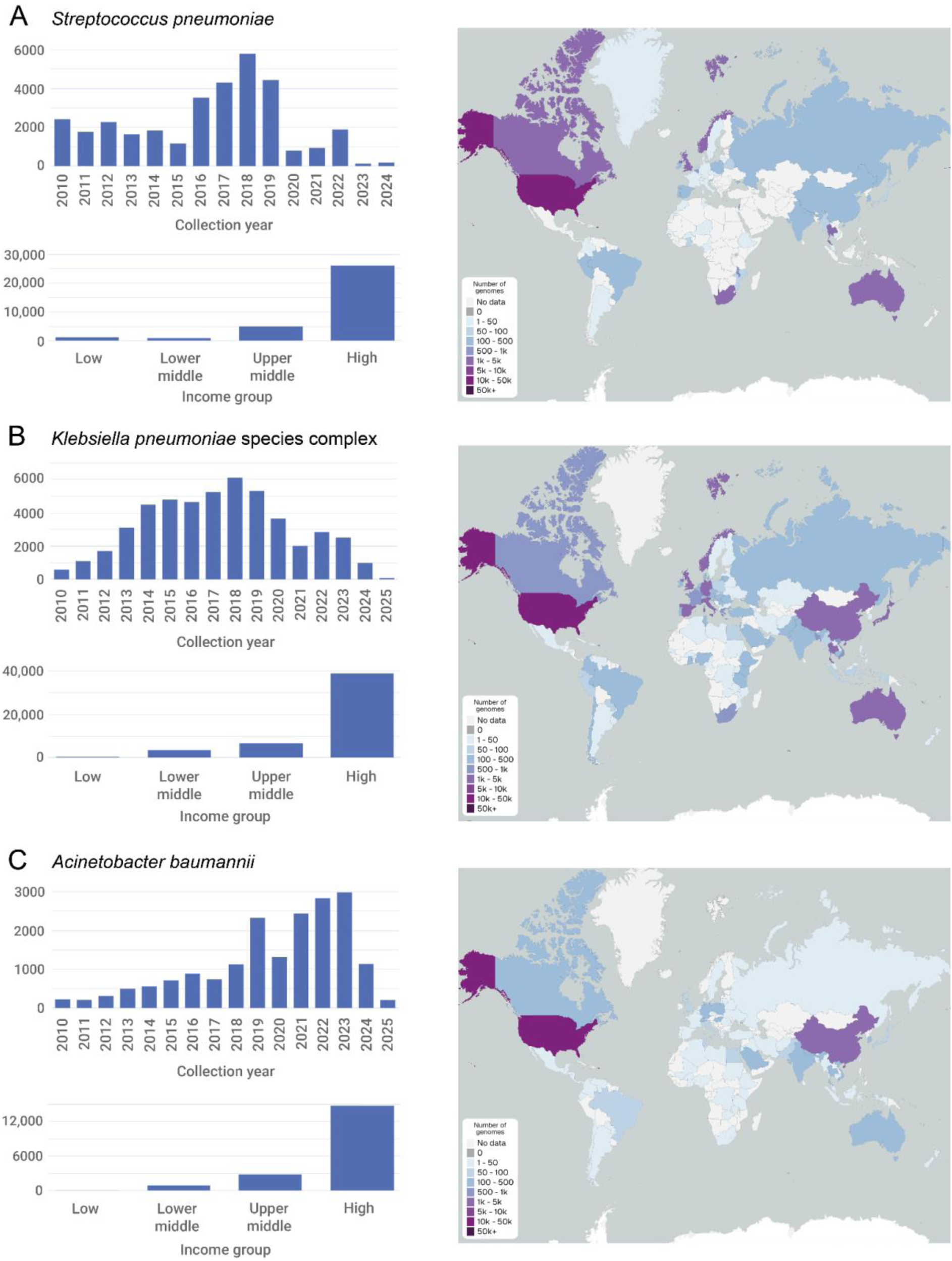
Distribution of genomes represented in vaccines.watch as of 3 June 2025. Panels show the number of genomes by collection year, by country income group and by country (map) for each of the three pathogens, *Streptococcus pneumoniae* (A), *Klebsiella pneumoniae* species complex (B) and *Acinetobacter baumannii* (C). A live overview of all genomes represented in vaccines.watch is available at https://vaccines.watch/all

The temporal distribution of genomes differed across the three pathogens (Figure 2). In particular, we found the majority of genomes from both *S. pneumoniae* (88.1%; 29,010/32,918) and *K. pneumoniae* SC (75.5%; 36,999/49,035) were sampled prior to 2020, with a decline observed since 2018. However, the number of *A. baumannii* genomes continued to increase until 2023, with the majority (59.0%; 10,865/18,428) sampled from 2020 onwards.

As detailed previously [13], we also found that high proportions of the public genomes carried one or more genes and/or mutations associated with AMR, albeit with the proportions differing considerably between countries. Overall, 32.7% (10,755/32,918) of *S. pneumoniae* genomes carried one or more AMR mechanisms associated with beta-lactam resistance. 74.3% (36,436/49,035) and 57.2% (28,033/49,035) of the *K. pneumoniae* SC genomes carried mechanisms associated with third-generation cephalosporin and carbapenem resistance, respectively. 83.5% (15,385/18,428) of the *A. baumannii* genomes carried mechanisms associated with carbapenems, which rose to >90% for some individual countries including Brazil (97.9%; 93/95), Vietnam (95.7%; 111/116) and Poland (95.3%; 143/150).

Altogether these findings continue to show the strong biases that exist among available public genomes, reflecting the varying availability of genome sequencing worldwide to date and its use within specific research agendas that often have a major focus on AMR.

### Assessing trends in vaccine target diversity from global genome data

We next reviewed the vaccine target diversity observed among the global public genomes in vaccines.watch (available as of 3 June 2025), albeit with awareness of the limitations relating to the representativeness of currently-available data. Using examples from across the three different pathogens, we illustrate how vaccines.watch can be used to interrogate trends among the vaccine targets in the context of available vaccines, their geotemporal dynamics and the broader population diversity.

#### Streptococcus pneumoniae

The vaccines.watch platform displays data on the predicted serotypes of *S. pneumoniae* based on identification of the corresponding *cps* loci by SeroBA. Among the 32,918 *S. pneumoniae* genomes, we found a total of 89 different serotypes from a total of 102 that can currently be identified by SeroBA. The top twenty most frequent serotypes accounted for 76.2% (25,075/32,918) of the total genomes while many serotypes were rare (e.g. 38 serotypes accounted for <20 genomes each). Only 0.1% (42/32,918) of genomes were untypeable by SeroBA, which can reflect either true absence of an intact *cps* locus or poor-quality data. A further 0.7% (219/32,918) of genomes had matches to “null capsule clade” (NCC) (i.e. non-encapsulated) variants.

As described above, vaccines.watch enables users to select specific pneumococcal vaccine formulations and review their composite serotypes among all (or selected) genomes. For example, we observed that serotypes included in the PCV13 formulation accounted for 36.2% (11,907/32,918) of the global public genomes. Among the highest-sampled countries, this proportion varied between 20.3% and 55.7% (USA (30.3%), South Africa (49.3%), UK (20.3%), Norway (28.3%), Australia (36.0%), Canada (35.7%), Thailand (55.7%)) (**Figure 3**). Serotypes included in the broadest vaccine to date, PCV31 (VAX-31), currently in clinical trials [29], accounted for 87.4% (28,764/32,918) of global genomes.

**Figure 3.**
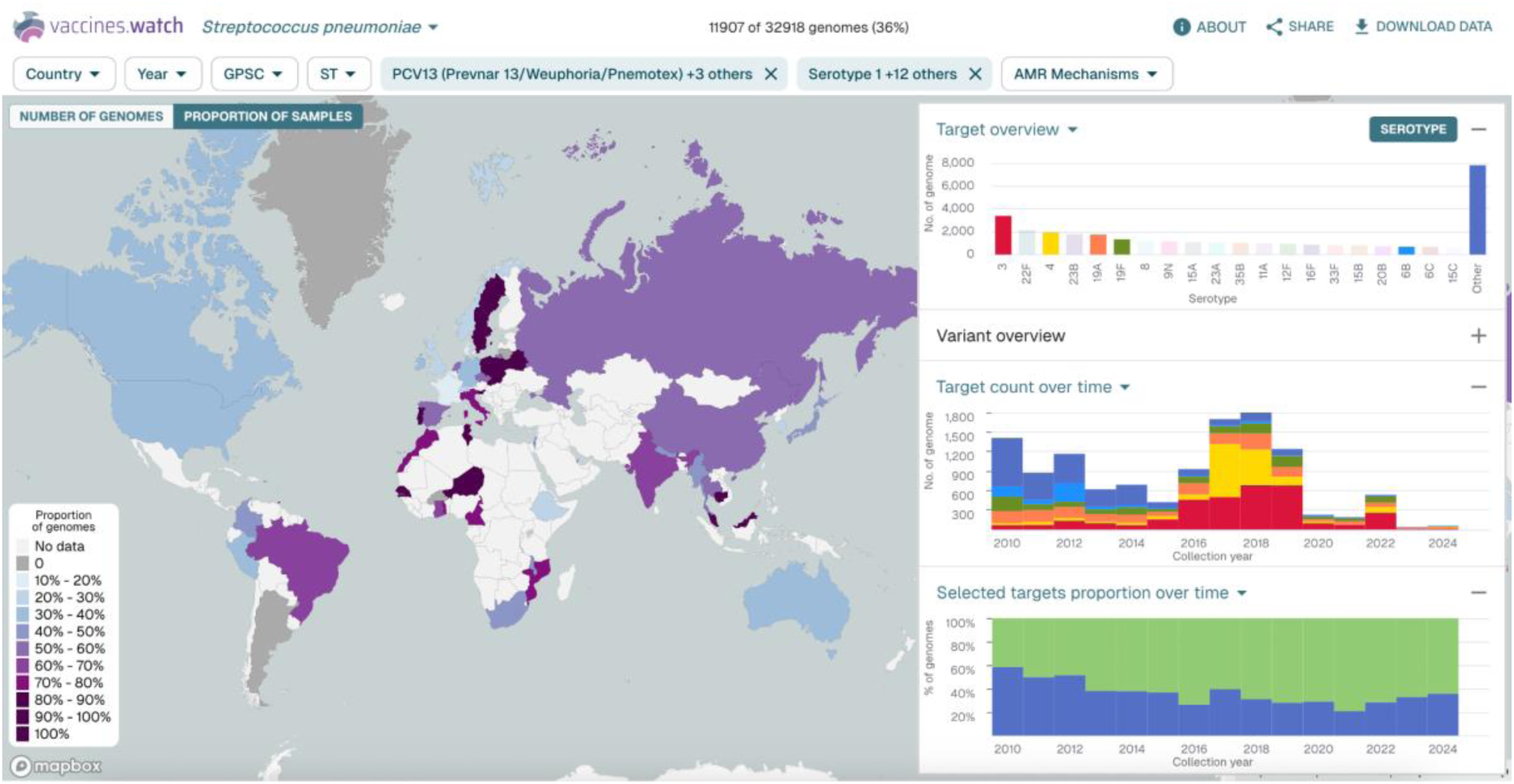
Assessment of serotypes targeted by the PCV13 pneumococcal vaccine formulation in the context of the global *S. pneumoniae* genomes included in vaccines.watch (as of 3 June 2025). The map shows the proportion of genomes in each country that belong to one of the 13 serotypes targeted by PCV13. The right-hand panels show the twenty most frequent *S. pneumoniae* serotypes with PCV13 serotypes highlighted (top), the distribution of PCV13 serotypes by sampling year (middle) and the proportion of the total genomes that belong (blue) or do not belong (green) to one of the PCV13 serotypes by sampling year (bottom). A similar visualisation can be found at: https://cgps.dev/qM98oI6

Despite its inclusion within PPSV23 and all PCV formulations since PCV13, serotype 3 was the most frequent serotype among all genomes, accounting for 10.3% (3400/32,918) of the total (Figure 3). While this may partially reflect the variable global coverage of pneumococcal vaccines, studies have also shown that this serotype has continued to be a major cause of invasive pneumococcal disease even within countries that have achieved high coverage of PCV13 such as Spain [30], with some evidence of low vaccine effectiveness against this serotype [31].

Vaccines.watch can also be used to identify serotypes that are prevalent among public genomes but not included in vaccine formulations (Figure 3). For example, we observed that 23B, which is not included in PPSV23 or early PCV formulations (e.g. PCV13), was the fourth most prevalent serotype among the public genomes (accounting for 5.5% (1795/32,918)). This serotype has been shown to be associated with increasing prevalence and penicillin non-susceptibility among both carriage and invasive isolates in recent years [32].

#### Klebsiella pneumoniae species complex

The vaccines.watch platform displays data from Kaptive on both the best-matching K/O locus types (genotypes) identified from the *K. pneumoniae* SC genomes and the predicted K/O types (phenotypes), using the new O serotyping nomenclature proposed recently [33]. We have included both genotypic and predicted phenotypic types to allow exploration of the corresponding relationships. In the case of the O loci/types, these do not conform to one-to-one relationships due to the use of both the O locus and additional genes from outside of the O locus in the phenotype predictions.

Among the 49,035 *K. pneumoniae* SC genomes, we identified 139 different K loci from the 186 that are currently defined by Kaptive. 3.2% (1592/49,035) of the total genomes were predicted to encode an acapsular phenotype (i.e. “capsule null”), based on identification of a K locus with truncations in one or more essential capsular synthesis genes. The top twenty most frequent K loci (without essential gene truncations) accounted for 64.7% (31,743/49,035) of the total genomes. Many of the K loci were found rarely (e.g. 20 accounted for <20 genomes each). Among genomes with carbapenem resistance markers, the top twenty K loci (without essential gene truncations) accounted for 79.3% (22,242/28,033). Notably, we also found that 35.0% (17,139/49,035) of all genomes carried a K locus that corresponds to an unknown K type. These K loci include some of the most frequent in the collection, such as KL107, KL102 and KL106, which accounted for the second, third and fifth highest number of genomes, respectively.

The diversity of O locus types found among the public genomes was lower, with a total of 12 identified. These correspond to 22 predicted O types (serotypes), the full repertoire currently defined by Kaptive. Five O types (O2β; O1ɑβ,2ɑ; O2ɑ; O1ɑβ,2β; O3γ) accounted for 80.3% (39,351/49,035) of the genomes. The two O locus types, OL2ɑ.1 or OL2ɑ.2, which underlie the different array of O1 and O2 types (together with OL2ɑ.3 which was found rarely), accounted for 72.0% (35,325/49,035) of all genomes.

Using vaccines.watch, we can observe that the distribution of both K and O antigens among public genomes varies by country (**Figure 4**). This is largely related to the variable geographic distribution of STs, which are often dominated by single K and O loci/types. Assessment of the distribution of K antigens over time showed a proportional increase in KL64 (encoding K64) from around 2014, with it now being the most prevalent K locus type, accounting for 22.5% (219/975) of genomes from 2024. This could be linked to an increase in ST147 genomes, 81.1% (2363/2913) of which possess KL64. We also noted a rise in O2ɑ since 2020, accounting for 38.4% (374/975) of genomes from 2024, and mostly encoded by the OL2 ɑ.1 variant. We could observe that this rise in O2ɑ has been driven by a proportional increase in genomes from both ST147 and ST45, 82.1% (2393/2913) and 92.4% (1112/1203) of which encode O2ɑ, respectively.

**Figure 4.**
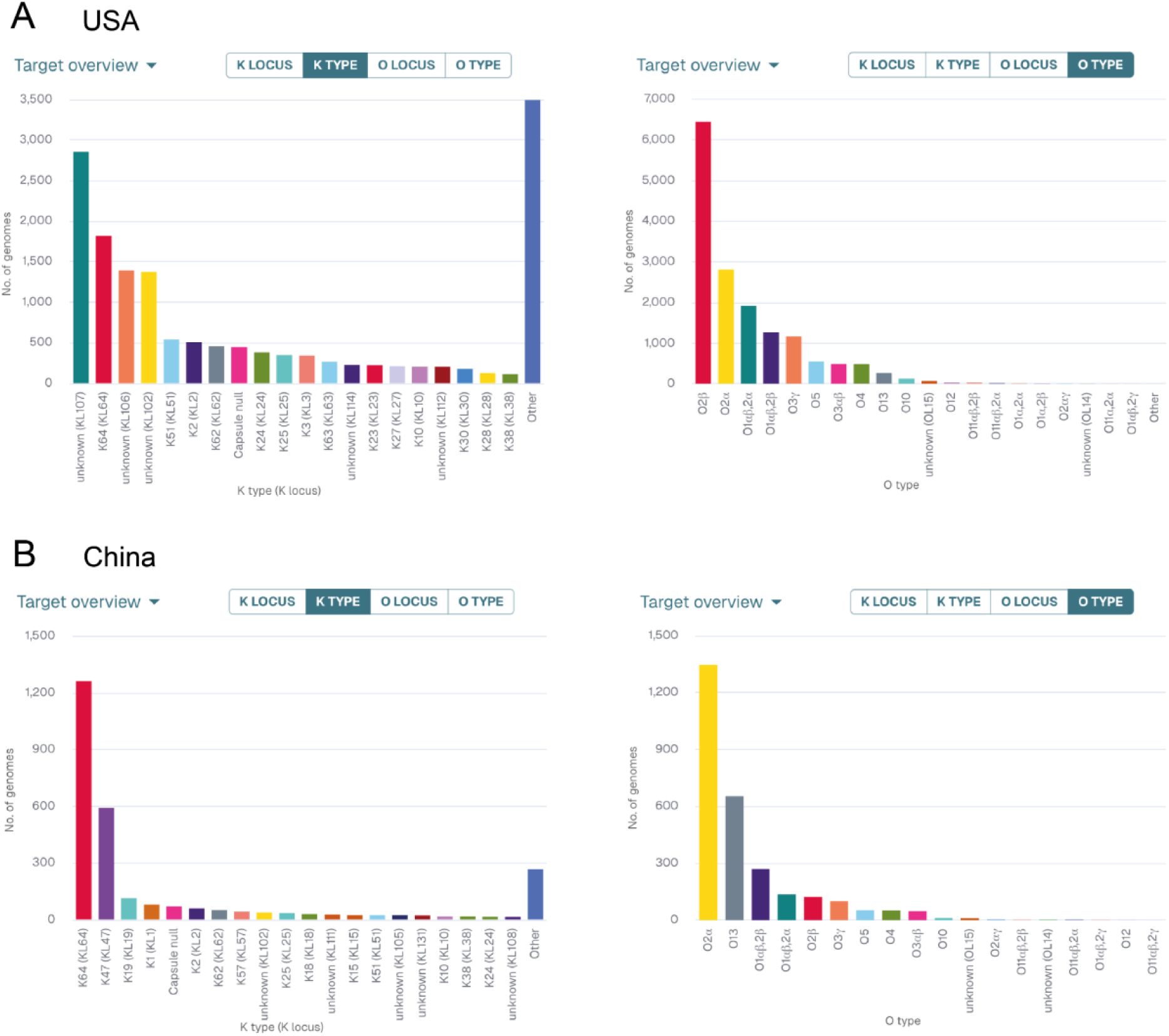
Number of genomes with predicted K and O types among *K. pneumoniae* species complex from the USA (A) and China (B), based on genomes represented in vaccines.watch as of 3 June 2025. Similar visualisations, updated in real-time, are available at: https://vaccines.watch/organism/570?Country+Code=US (A) and https://vaccines.watch/organism/570?Country+Code=CN (B).

**Figure 5.**
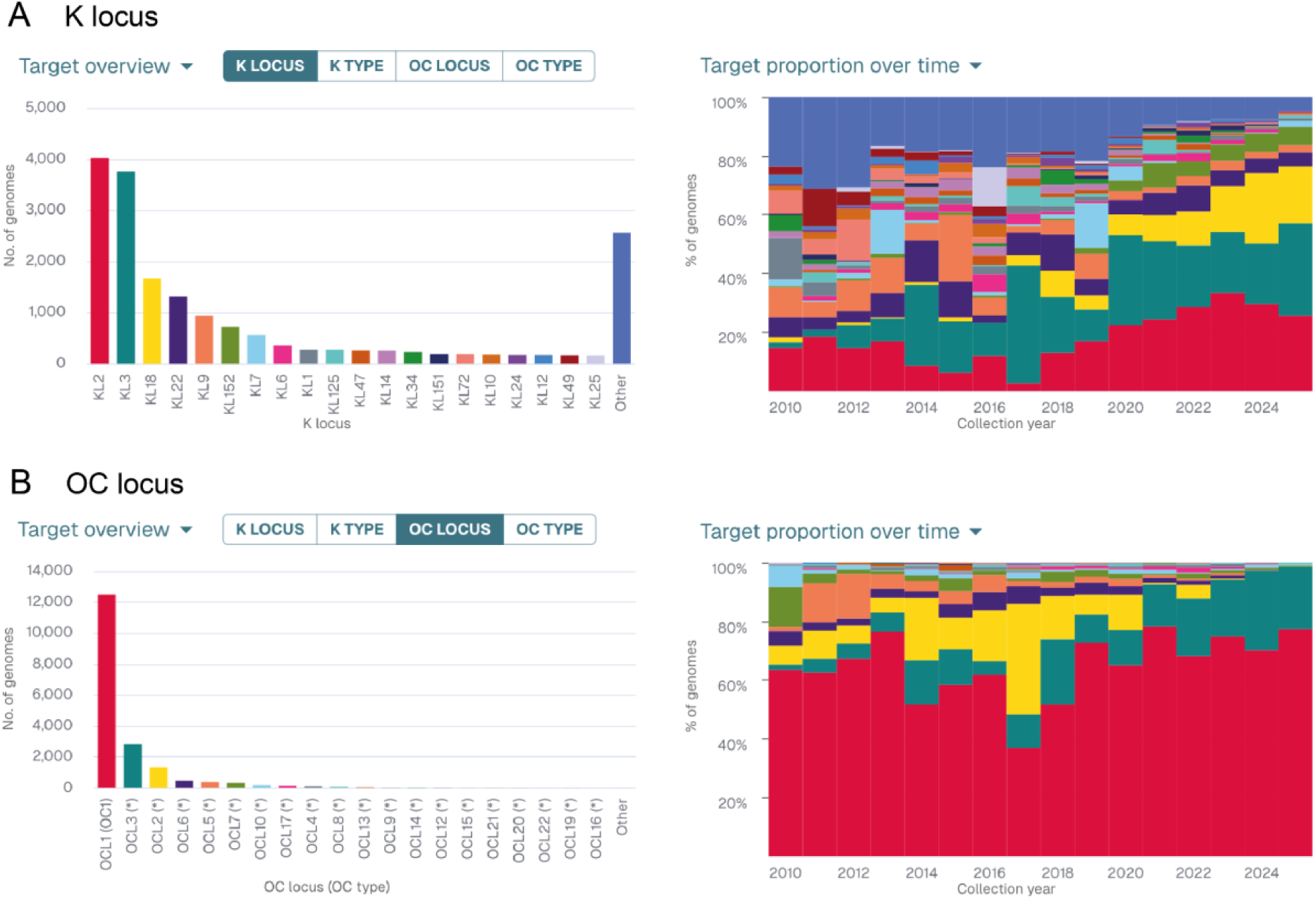
Distribution of K (A) and OC (B) locus types among *A. baumannii*, based on genomes represented in vaccines.watch as of 3 June 2025. The plots show the number of genomes with different locus types, ordered by frequency (left), and the proportion of the different locus types by sampling year, using the same colouring of types (right). Similar visualisations, updated in real-time, can be found at https://vaccines.watch/organism/470

#### Acinetobacter baumannii

Similarly as for *K. pneumoniae* SC, vaccines.watch displays data from Kaptive on both the best-matching K/OC locus types (genotypes) identified from the *A. baumannii* genomes and the predicted K/OC types (phenotypes). As with the O loci/types in *K. pneumoniae* SC, the K loci/types in *A. baumannii* also do not conform to one-to-one relationships due to the use of both the K locus and additional genes in the phenotype predictions. Notably, the *A. baumannii* genome collection is highly dominated by a single variant type, ST2, from an internationally-distributed lineage known as global clone 2 (GC2) [34], which comprises 64.4% (11,866/18,428) of the total genomes.

We investigated the diversity of K loci and predicted types, which are known to be the major immunodominant antigens in *A. baumannii* (16, 35], as well as the primary determinants of phage susceptibility [36]. We found 194 K loci among the 18,428 *A. baumannii* genomes, from a total of 241 currently defined by Kaptive. Despite the high overall diversity, KL2 (encoding K2) and KL3 (encoding K3 and K3-v1) dominated the public genome collection, accounting for 21.8% (4018/18,428) and 20.4% (3757/18,428) of the total genomes, respectively. The top twenty K locus types together accounted for 86.1% (15,867/18,428) of the total genomes, while 141 of the KL types identified were rare (present in <20 genomes). Of the genomes possessing KL2, 96.6% (3883/4018) were from ST2. Among those with KL3, 67.4% (2533/3757) were from ST2 while a further 22.9% (859/3757) were from ST437. Notably, we also found that 23.5% (4323/18,428) of all genomes possessed a K locus that corresponds to an unknown K type.

Assessment of the diversity of OC loci identified 22 locus types among the public genomes, corresponding to all of the known OC locus types defined within the Kaptive database. However, the genomes were highly dominated by OCL1 (encoding OC1) which accounted for 67.8% (12,499/18,428), with OCL3 accounting for a further 15.3% (2816/18,428) and OCL2 accounting for 7.3% (1341/18,428). 87.4% (10,919/12,499) of genomes with OCL1 were from ST2.

Among both K locus and OC locus types, we observed a general trend towards lower diversity over time (**FIgure 5**), which may reflect increased sampling of major multidrug-resistant lineages in recent years. In particular, we observed an increase in both KL18 and OCL3 from 2020 onwards. This could largely be attributed to a rise in the proportion of ST499, which accounted for 84.2% (1408/1672) and 50.1% (1412/2816) of genomes with KL18 and OCL3, respectively. ST499 is reported to have emerged as a dominant non-GC2 carbapenem-resistant lineage in the USA in recent years [37]. ST499 genomes were primarily obtained from isolates sampled in the USA (99.2%; 1400/1412), with a further nine genomes from Fiji and a single genome obtained from each of Canada, Germany and India.

## Discussion

As pathogen genome sequencing becomes increasingly integrated into routine disease surveillance, there are growing opportunities for the shared global data to enhance precision around the targeting of public health interventions. Here we present an interactive platform, vaccines.watch (https://vaccines.watch), which aims to guide efforts in vaccine research and development by enabling interrogation of vaccine target diversity from genome data within the context of the pathogen population dynamics. The platform currently displays data from high-quality public genomes with geotemporal sampling information (post-2010), which are incorporated on an ongoing basis as genomes are deposited in INSDC databases. We have initially focused on developing vaccines.watch for polysaccharide targets from major bacterial pathogens which form the basis of existing or prospective multivalent vaccines. In particular, we have included the capsular polysaccharide targets from pneumococcal vaccines for which there remains an ongoing need for improved and/or updated formulations despite their high efficacy in reducing invasive pneumococcal disease [38]. We have also included capsular and LPS O antigen-based targets from *K. pneumoniae* SC and capsular and LOS outer core-based targets from *A. baumannii*. Both *K. pneumoniae* and *A. baumannii* are leading multidrug-resistant pathogens that pose a critical public health threat [39, 40], particularly to vulnerable patients (including neonates) within hospital settings. There is only a limited vaccine pipeline for *K. pneumoniae* (one O antigen-based candidate, Kleb4V, completed a phase 1/2 trial in 2022 (NCT04959344)) [41] while there are currently no vaccines in active clinical development for *A. baumannii* [4].

As the global representativeness of genome data grows, we anticipate that vaccines.watch will aid the selection of vaccine targets for new or updated multivalent vaccine formulations. The choice of which precise target types to include within vaccine formulations, such as the *S. pneumoniae* serotypes within pneumococcal vaccines, is critical due to factors such as the varying frequency among target populations, invasiveness potential and antimicrobial resistance profiles associated with different types [42, 43]. Due to geographic differences among pathogen populations, we envisage that new vaccine formulations will be increasingly tailored to specific localities to enable maximal protection against disease. For example, a 10- valent PCV (Pneumosil) has already been designed to provide protection against serotypes causing the highest disease burden in Africa, Asia, Latin America and the Caribbean [44]. Increasing availability of representative genomic data may also be used to assess the likely efficacy of existing vaccine formulations within different localities and/or target populations, and inform choice where multiple vaccines with differing compositions exist. This approach has the potential to expand the use of existing vaccines, including pneumococcal vaccines which still have highly variable coverage worldwide [45].

We also anticipate that, with improving genome representation, vaccines.watch could be used for monitoring the impact of new vaccines by enabling comparison of target types and the associated population diversity before and after vaccine introduction. Post-vaccination monitoring is especially critical with polysaccharide-based vaccines, as these target antigens are known to exhibit high rates of recombinational exchange and therefore are liable to “switching” [46, 47]. Numerous studies have demonstrated the value of genomic data for high-resolution monitoring of vaccine-related population changes that, for example, have led to an increase in non-vaccine pneumococcal serotypes with higher invasiveness and/or AMR [48, 49]. In the case of pneumococcal disease, the ongoing approach to address these changes is to develop new PCV formulations with increasing serotype coverage, with licensed vaccines now including up to 21 serotypes [15].

The opportunities described above have the potential to launch an exciting new era of vaccine development yet remain highly dependent on global efforts to increase and sustain pathogen genome sequencing capacity for routine surveillance. Our review of global public genomes, which represent the current data source for vaccines.watch, highlights the substantial gaps in geographic coverage, paucity of recent data (likely, in part, due to a lag in deposition times), and ongoing biases in sequencing which remains dominated by specific research agendas (e.g. relating to AMR). We therefore advise users to remain highly vigilant to these data limitations and maintain careful consideration of how data shown in vaccines.watch is used and interpreted. Another important limitation in the current use of genomic data for vaccine development and monitoring efforts is that there is still an incomplete understanding of how the genomic information relates to the polysaccharide phenotypes, with many genomic loci encoding unknown structures [33], including across all three pathogens currently included within vaccines.watch. However, for each of these three pathogens, there are active efforts to improve this knowledge base, with regular updates made to the nomenclature databases and phenotype prediction logic provided by the SeroBA [27] and Kaptive tools [25]. In particular, for *S. pneumoniae*, there is a comprehensive and curated library, SeroBAnk, collating data on the genetic locus and capsular structure of each known serotype [27].

Future developments of vaccines.watch include extending our approach to additional vaccine targets, starting with polysaccharide targets from other bacterial pathogens with *in silico* typing schemes and that form the basis of similar multivalent vaccine formulations. We also aim to develop the ability to display data from bespoke genome collections that may be pre-defined within vaccines.watch or sourced by users themselves. Finally, we can readily adapt vaccines.watch to include other data types, such as additional metadata associated with pathogen genomes (e.g. source, disease type, patient characteristics). This will be important as available metadata becomes increasingly harmonised for individual pathogens, driven by recent ongoing curation efforts of the public health and research communities and development of metadata templates.

In conclusion we anticipate that, as genomic surveillance efforts increase, the use of large-scale genomic data will become an essential component of the vaccine developer’s toolkit. With the development of vaccines.watch, we have demonstrated how global genome data can be ingested, analysed and delivered in real-time via an accessible platform to provide insights into the pathogen population dynamics around key vaccine targets. The platform aims to support scientists and vaccine developers with decision-making around vaccine formulations and roll-out.

## Acknowledgements

We would like to thank Joshua Wong for helpful discussions regarding the development of vaccines.watch.

## Data availability

All data represented in vaccines.watch are available for download within the application. The assembled genomes and associated metadata can be accessed via Pathogenwatch (https://next.pathogen.watch), together with additional genotypic data. Raw sequence reads are available in the ENA/SRA.

